# Co-circulation of distinct high pathogenicity avian influenza virus (HPAIV) subtypes in a mass mortality event in wild seabirds and co-location with dead seals

**DOI:** 10.1101/2025.07.11.664278

**Authors:** Marco Falchieri, Eleanor Bentley, Holly A. Coombes, Benjamin C. Mollett, Jacob Terrey, Samantha Holland, Edward Stubbings, Natalie Mcginn, Jayne Cooper, Samira Ahmad, Jonathan Lewis, Ben Clifton, Nick Collinson, James Aegerter, Divya Venkatesh, Debbie J. F. Russell, Joe James, Scott M. Reid, Ashley C. Banyard

## Abstract

H5Nx clade 2.3.4.4b high pathogenicity avian influenza viruses (HPAIV) have been detected repeatedly in Great Britain (GB) since autumn 2020, with H5N1 dominating detections but with low level detection of H5N5 during 2025. Globally, these viruses have caused mass mortalities in captive and wild avian and mammalian populations, including terrestrial and marine mammals. H5N1 has been the dominant subtype, and whilst incursions have overlapped temporally, occurrences have often been spatially distinct. Here, we report the detection of a mortality event in wild birds on the Norfolk coastline in the east of England, where H5N1 HPAIV was detected in five Great Black-backed Gulls (*Larus marinus*) and a Northern Fulmar (*Fulmarus glacialis*). Interestingly, at the same site, and as part of the same mortality event, a total of 17 Great Black-backed Gulls, one Herring Gull (*Larus argentatus*), one Atlantic Puffin (*Fratercula arctica*) and one Northern Fulmar tested positive for H5N5 HPAIV. Additionally, H5N5 was also detected in 17 co-located Grey Seal carcases (*Halichoerus grypus)*. The H5N1 HPAIV from an infected bird belonged to genotype DI.2, closely related to contemporaneous detections in GB wild birds and poultry. In contrast, all H5N5 HPAIVs from birds and seals were genotype I with a 22-amino acid stalk deletion in NA and the 627K polymorphism in PB2. This represents the first recorded instance in GB of two subtypes being detected within the same avian population at the same location. It is also the first mass detection of HPAIV H5N5 in mammals within GB. Potential infection mechanisms are discussed.

## Introduction

H5Nx clade 2.3.4.4b high pathogenicity avian influenza viruses (HPAIVs) have been consistently detected in Great Britain (GB) since autumn 2020, affecting both wildlife (avian and mammalian species) and poultry farms [1, 2]. The early phase of the outbreak (autumn 2020 to spring 2021) was dominated by the H5N8 HPAIV subtype, while H5N1 HPAIV has been the dominant viral subtype detected since summer 2021 [3, 4]. The dominance of the H5N1 subtype has been seen globally with mass mortality events in wild and captive birds and sporadic spillover events into a limited number of wild, free-living mammals [2, 5]. Additionally, H5N5 HPAIV has been intermittently detected since autumn 2023 in GB, primarily affecting avian species as well as being sporadically detected across Europe [6], North America [7] and Asia [8]. In Europe, detection in wild birds is often found with wild species associated with marine and coastal contexts. Interestingly, from an avian perspective, whilst the finding of the H5N1 and H5N5 HPAIV subtypes have overlapped temporally, their detections have typically been ecologically distinct.

The detection of mass mortality events across multiple farmed and wild mammals is a recent feature of outbreaks with a virus that has previously only affected avian species, but the scale of the epizootic in reservoir species seem to be the main cause of these spill-over events where distribution and ecologies overlap. This is particularly true of outbreaks in marine mammals, such as those seen in South America, where a range of pinniped species (carnivorous aquatic mammals such as seals) have been involved [9-11] and genetic assessment of viruses has demonstrated the interplay of the virus between avian and mammalian hosts. This has also been demonstrated in sub-Antarctic environments where mortality in birds has led to mortalities in Antarctic Fur Seals (*Arctocephalus* gazella) and Southern Elephant Seals (*Mirounga leonina*) [12, 13]. A further dimension has been observed in farmed mammalian species including mink [14], foxes [15], sheep [16], and most notably dairy cattle (in the United States of America) [17]. The latter has increased the zoonotic potential of this virus and pandemic risks through the excretion of very high titres of virus in milk from infected dairy cattle [18-20]. Not only has this led to increased risk of human exposures to the virus, albeit generally with mild infection outcomes, but both wild and domestic avian and mammalian species have also succumbed, likely through exposure to infected milk [17]. A further risk to animal health has come through environments where captive (e.g., zoological collections) [21] or domesticated animals (e.g., pet cats) have been fed contaminated avian material as foodstuffs [22]. Although comparatively rare, this route of exposure has been reported on numerous occasions with significant outcomes as animals have succumbed to infection via this route. From a GB perspective, in wild free-living mammals, sporadic detections of infection with HPAIV H5Nx have been detected [23] but no mass mortality events have been described.

Influenza A virus infections of seals are poorly understood and are only sporadically detected, with other pathogens generally being responsible for mortality events. There are two species of seals that live in the British Isles, Grey Seal (*Halichoerus grypus*), and Common Seal or Harbour Seal (*Phoca vitulina*), but seal populations vary in size and location. One key area where seal populations visit and are monitored in GB is Blakeney Point (52.9752°N, 0.9921°E), Norfolk [24]. Around 10% (>7,400 pups) of GB grey seal pups are born at Blakeney Point, and it is the largest seal breeding colony in England [25]. The GB breeding figures account for ∼35% of the global and ∼90% of the North-east Atlantic populations. [25]. During the breeding season at Blakeney (November to January), females congregate to give birth to a single pup which they suckle for ∼18 days before pups moult their white coats (lanugo) and take to the sea at ∼31.5 days. The females come into oestrus at weaning, attracting the presence of males. The breeding season is asynchronous so not all seals will be at the same location at the same time but nevertheless, ∼20,000 Grey Seals (males, females and pups) likely visit Blakeney Point during the breeding season [24]. This substantial and dense aggregation suggests the potential for two-way transmission of disease between seals and birds (mainly gulls), with likely exposure to seabird faeces and carcases on the shoreline and seal haul out, and the scavenging by birds of seal afterbirths and dead pups by gulls [26, 27]. There is likely exposure to seabird faeces, as well as to dead birds, in the pupping areas where large gulls (Herring Gull and Great Black-backed Gulls) feed on dead seals and afterbirths. Other birds such as the Puffin and Fulmars had likely been washed in by the sea, but infected dead bodies could still provide an interface for pathogen exchange.

Here, we report the occurrence of a mass mortality event in wild birds in the East of England, where both H5N5 HPAIV and H5N1 HPAIV were detected in gulls (members of the order Charadriiformes, family Laridae), alongside concomitant infection of Grey Seals with H5N5. This represents the first recorded instance in the UK of H5N1 and H5N5 being detected as co-circulating at the same location and within the same species, as well as the first detection of HPAIV H5N5 in mammals. We overview the case history, clinical manifestations, and epidemiological investigation, along with virological, molecular, and bioinformatic analyses that confirm these findings and hypothesise infection routes and the potential directionality of infection based on details of the incident.

## Materials and Methods

### Clinical and epidemiological investigation

An avian mortality event investigation was triggered following a report to the Department for Environment, Food & Rural Affairs (DEFRA) online reporting system [28] of bird carcasses on National Trust land at Blakeney Point, Norfolk. This is a relatively remote extensive dune and sandy beach coast which is managed as a nature reserve and is protected by numerous official designations. The foreshore is wide and long, with numerous sandbars on which seals haul out and breed. This part of the coast forms the southern edge of ‘The Wash’, a geographically extensive combination of sea bay, coastal and estuarine habitats which together hosts the single largest aggregation of wintering waterbirds in the UK. Samples from bird carcasses were taken in line with government guidelines to determine wild bird species affected, their number, the level of environmental contamination, as well as the distribution of viral subtypes involved. Seal carcasses were also observed during the preliminary field visit, at a time where a natural baseline of ∼5 % mortality was expected [29]. Considering recently reported HPAIV H5Nx cases in pinnipeds across the globe and the potential for mass mortality events potentially including mammal-to-mammal transmission, a further epidemiological investigation was carried out to assess whether any of the dead seal mortalities were positive for influenza virus as well as to determine the range of avian species affected, the level of environmental contamination, and the distribution of HPAIV subtypes involved.

Multiple samples were collected from a range of carcasses (avian and mammalian) as well as avian faecal material (n=8), feathers (n=3) and a water sample (n=1) as part of the environmental and virological assessment. Samples from carcasses included swabs from orifices (Oral, Nasal and Rectal for mammals, and oropharyngeal (OP) and cloacal (C) for birds) and samples from target internal organs (either swabs or tissues), such as brain, lung and miscellaneous.

### Virological investigation

Sample processing depended on sample type and matrix and were processed as described previously [30]. Swabs samples were cut into 1 mL serum-free Leibovitz’s L-15 medium (Gibco) containing antibiotics (penicillin, streptomycin), incubated at room temperature for 10 min before standard viral RNA (vRNA) extraction. Brain and lung tissues were prepared as a 10% (w/v) suspension in L-15 medium and incubated at room temperature for 60 min before using standard RNA extraction protocols [31]. RNA was extracted using the MagMAX CORE Nucleic Acid Purification Kit (ThermoFisher Scientific) as previously described [31]. Extracted RNA was assessed for vRNA using RT-PCR assays specific for M gene [32], an H5 HPAIV [33] and/or NA [34]. A standard curve was generated using a 10-fold dilution series of titrated H5N1 HPAIV RNA as previously described to assess test efficiency [31].

### Genomic analysis

Whole genome sequences were generated from positive samples using Oxford Nanopore Technology as described previously [12]. Briefly, extracted vRNA was converted to double stranded cDNA and amplified using a one-step RT-PCR using SuperScript III One-Step RT-PCR kit (Thermo Fisher Scientific), see Supplementary Table 1 for specific primers. PCR products were purified with Agencourt AMPure XP beads (Beckman Coulter) prior to sequencing library preparation using the Native Barcoding Kit (Oxford Nanopore Technologies) and sequenced using a GridION Mk1 (Oxford Nanopore Technologies), according to manufacturer’s instructions.

Assembly of the influenza A viral genomes was undertaken using a custom in-house pipeline previously described [35]. Comparison of the study-derived sequences and contemporary H5 sequences was undertaken against all avian H5 sequences available on GISAID between the 1^st^ of January 2020 and the 28^th^ of February 2025. All sequences were aligned on a per segment basis using MAFFT v7.520 [36] and trimmed against a reference using SeqKit v2.5.1 [37]. The trimmed alignments were used to a infer maximum-likelihood phylogenetic trees using IQ-Tree version 2.2.5 [38] along with ModelFinder [39] and 1000 ultrafast bootstraps [40]. Ancestral sequence reconstruction and inference of molecular-clock phylogenies were undertaken using TreeTime v0.10.1 [41]. Phylogenetic trees were visualised using R version 4.3.3 with ggplot2 version 3.5.1 [42] and ggtree version 3.14.0 [43]. Sequences were genotyped from phylogenetic trees by comparison to known reference sequences for all genotypes currently circulating in GB. Sequences derived from both seals and avian species were assessed for the presence of adaptive mutations that may confer increased replication in mammals. All sequences were aligned on a per segment basis using MAFFT v7.520 [36] and manually trimmed to the open reading frame using Aliview version 1.28 [44]. Trimmed sequences were translated to amino acids and visually inspected for mutations. All influenza sequences generated in this study are available through the GISAID EpiFlu Database (https://www.gisaid.org, Supplementary Table 2)

## Results

### Clinical and epidemiological investigation

An investigation into a mortality event involving gulls and other avian species was initiated at Blakeney Point, Norfolk (Figure 1), on the 5^th^ of February 2025 under the avian influenza wild bird surveillance scheme [28]. Carcasses were in varied states of decomposition and often incomplete, sometimes discovered as disarticulated limbs, heads etc. In addition, the beach was extensive and a complete search in the time available between tides was impossible. As such, a precise carcass count could not be defined but the likely number of dead birds far exceeded the number sampled. In total, five birds including four Great Black-backed Gulls (GBBG; *Laurus marinus*) and one Herring Gull (*Larus argentatus*) were identified in the field and sampled; four Grey Seal carcasses were also sampled (Table 1 and supplementary Table 3).

**Figure 1:**
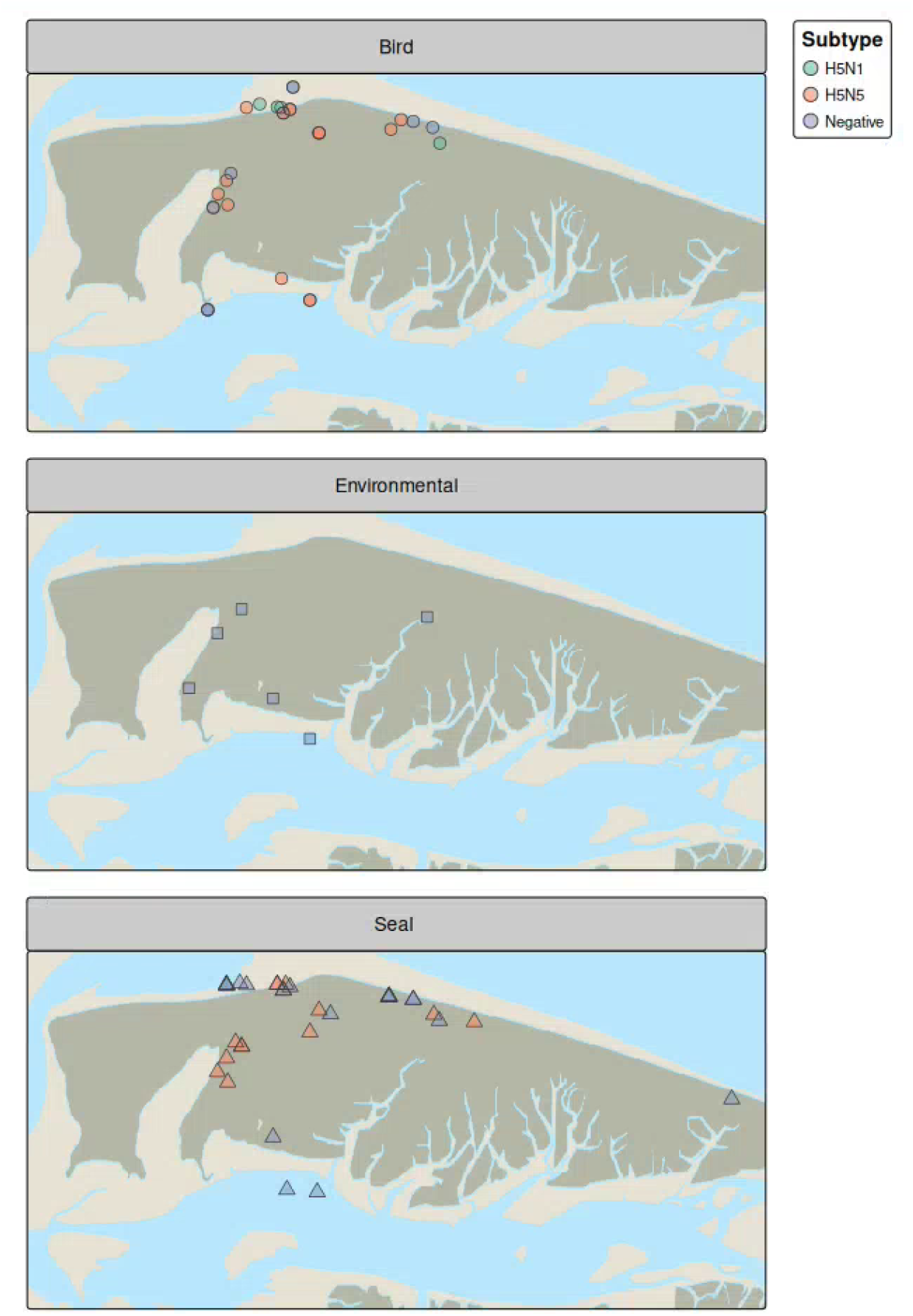
Spatial representation of samples collected from birds, the environment and seals and H5N1 or H5N5 HPAIV testing results.

**Table 1:**
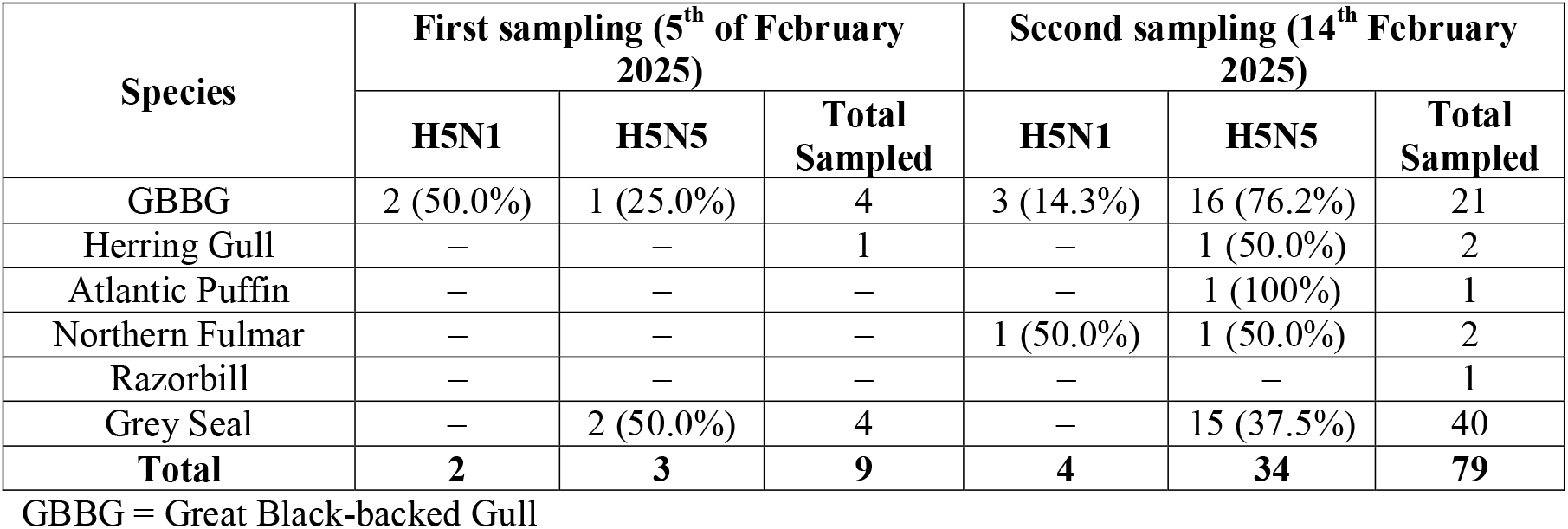
summary of animals sampled and HPAIV H5N1 and H5N5 positive animals during the incident.

| Species | First sampling (5<sup>th</sup> of February 2025) |  |  | Second sampling (14<sup>th</sup> February 2025) |  |  |
| --- | --- | --- | --- | --- | --- | --- |
|  | H5N1 | H5N5 | Total Sampled | H5N1 | H5N5 | Total Sampled |
| GBBG | 2 (50.0%) | 1 (25.0%) | 4 | 3 (14.3%) | 16 (76.2%) | 21 |
| Herring Gull | – | – | 1 | – | 1 (50.0%) | 2 |
| Atlantic Puffin | – | – | – | – | 1 (100%) | 1 |
| Northern Fulmar | – | – | – | 1 (50.0%) | 1 (50.0%) | 2 |
| Razorbill | – | – | – | – | – | 1 |
| Grey Seal | – | 2 (50.0%) | 4 | – | 15 (37.5%) | 40 |
| **Total** | **2** | **3** | **9** | **4** | **34** | **79** |

GBBG = Great Black-backed Gull

The results from these samples prompted a secondary sampling undertaken on the 14^th^ February 2025, whereby a total of 40 Grey Seal carcasses were sampled as well as 27 avian carcasses of various species including 21 GBBGs, two Herring Gulls, one Atlantic Puffin (*Fratercula arctica*), two Northern Fulmars (*Fulmarus glacialis*), and a Razorbill (*Alca torda*) (Table 1 and supplementary Table 3).

Mortality counts carried out by on-site staff at the end of January and beginning of February 2025 showed that an unusual die-off had taken place in Laridae with 46 GBBGs (both juveniles and adults) plus one Herring Gull and one Common Gull (*Larus canus*) being noted. However, the exact timing of the die-off is undefined as mortality counts are only performed at the end of the seal breeding season with timelines spanning from November 2024 up to February 2025. Most avian carcasses showed degrees of decomposition, mummification and scavenging, indicating that the avian die off had probably been ongoing throughout the winter. Blakeney Point, is home to England’s largest Grey Seal colony [24]. The site hosts and is visited by numerous wild birds including Anatidae (ducks, geese, and swans) and Laridae species. No large mortality events were recorded in seal pups during the 2024-2025 breeding season. Seal carcass counts undertaken between the 31^st^ of January 2025 and the 3^rd^ of February reported 177 seal carcasses present on site with only eight carcasses being adults and the remaining 169 carcasses constituting pups, weaners or immature seals. These mortality levels were within an expected mortality range for the colony during the breeding season. Further surveys, conducted after the 3^rd^ of February, estimated a total of up to 15 dead adults and 200 dead pups, including weaners and immatures.

### Virological and environmental investigations

Upon initial sampling 50% (n=2/4) Grey Seals and 60 % of avian sampled carcasses (n=3/5) tested positive for HPAIV, the latter including two GBBGs infected with HP H5N1 and one GBBG positive for HPAIV H5N5 (Table 1 and Figure 1). Following a second sampling effort, 37.5% (n=15/40) Grey Seals tested positive by PCR for HPAIV H5N5; no pinnipeds were positive for H5N1 HPAI. From an avian perspective, for the second sampling, a total of 27 birds were sampled: 76.2% (n=16/21) GBBGs, 50% (n=1/2) Herring Gull, 100% (n=1) Atlantic Puffin and 50% (n=1/2) Northern Fulmars tested positive for H5N5 HPAI; 14.3% (n=3/21) GBBGs and 50% (n=1/2) Northern Fulmar tested positive for H5N1 HPAI (Table 1 and Figure 1). Results per bird and sample type are shown in Supplementary Table 3.

### Genomic analysis

Eleven H5N5 HPAIV whole genome sequences (WGS) were recovered from a total of sixteen H5N5 clinical samples that were sent for sequencing (Supplementary Table 2). Six WGS were recovered from seals and one Herring Gull, one GBBG and one fulmar, with all seal- and avian-derived viral sequencing being H5N5 genotype I according to the EURL classification (Supplementary Figure 1). Mutational analysis demonstrated the presence of a 22-amino acid neuraminidase (NA) stalk deletion and a PB2 E627K polymorphism in all H5N5 samples.

One WGS was recovered from a GBBG, from a total of four H5N1 clinical samples that were sent for sequencing. This WGS belonged to the DI.2 genotype which has been the most prevalent in wild birds in the UK during winter 2024-2025. No mutations suggestive increased mammalian adaptation or increased zoonotic potential were found in the H5N1 sequence.

## Discussion

Incursions of HPAIV clade 2.3.4.4b H5Nx viruses into GB and Europe through migratory bird movements have occurred repeatedly since October 2020. The high-risk season for HPAIV incursion in GB typically starts each October coinciding with the beginning of the arrival of migratory wildfowl. This period also sees distinct behavioural changes in resident wildfowl which begin to aggregate on their overwintering grounds, often the same locations used by overwintering migrants. These changes in the size and contact structures of overwintering wildfowl promote the circulation and spread of HPAIV. Occasional spill-over events into commercial poultry and collections of kept bird collection can result in severe outbreaks of notifiable disease. Such outbreaks are controlled through rapid reporting, diagnostic testing, and movement restrictions, with culling of affected flocks to prevention further spread. As well as wild bird activity leading to infection of poultry, the reinfection of wild bird species from infected poultry has also been identified [45], although the frequency of bidirectional transmission is poorly understood. Significant mortality has also been seen in populations of wild birds [46-48], occasionally affecting species of conservation concern. Concern arises when notable disease events occur in nature reserves designated for the protection of vulnerable populations of overwintering waterbirds, such as Blakeney Point. High levels of HPAIV driven wild bird mortality increase infection pressure for mammalian species, especially scavengers.

The first mammalian wildlife detections of H5Nx in GB occurred in captive Common Seals, Grey Seals and a Red Fox (*Vulpes vulpes*) at a wildlife rehabilitation centre [49]. Diagnostic investigations linked the infections to a mortality event in Mute Swans (*Cygnus olor*) at the centre in the weeks preceding the detection in mammals. This initial detection involved the clade 2.3.4.4b H5N8 subtype and mammalian adaptations linked to increased zoonotic risk within viral sequences were limited to E627K and D701N within the PB2 gene [49]. Since 2021, clade 2.3.4.4b H5N1 has emerged as the dominant subtype across GB, Europe and the rest of the world with a vast array of genotypes being detected as part of that clade of H5Nx viruses [35, 50]. More recently, global detections of infection in different mammalian carnivore species [3, 4, 9-15] as well as herbivore/omnivore species [16, 17, 51, 52] have further underlined the ability of these viruses to cross the species barrier. Most recently, in GB, detection of H5N1 HPAIV (genotype DI.2), was confirmed in a single sheep, demonstrating risk across different sectors where infection pressure is high [16].

Although the initial clade 2.3.4.4b epizootic was caused by the H5N8 subtype, it was subsequently displaced by H5N1, which went on to dominate the global avian influenza landscape. Alongside, a further subtype, H5N5, has also been detected in different environmental contexts [7, 8]. In GB, HP H5N5 (genotype I) was first detected in Autumn 2023 with detections mainly in birds of prey and gull species. H5N5 was detected in 131 found dead wild birds, and one poultry premises between October 2024 up to the 28^th^ of February 2025 (through the statutory surveillance in wild birds) [53]. Until the event described here, H5N5 had never been detected in non-avian species in the British Isles.

The detection of H5N5 in both seals and gulls at Blakeney Point raises several questions regarding the infection and transmission kinetics of these viruses. The initial reason for sampling was a mortality event in gulls and in the weeks preceding this event, HP H5N1 wild bird positives were detected in the area predominantly associated with infection of Anatidae, birds of prey and gulls. All of the earlier detections were of the H5N1 DI.2 genotype that had become dominant both within the EU and across GB [54]. No local detections of H5N5 had been made until this event was investigated. It is not clear why a few gulls tested positive for H5N1 whilst most gulls and other avian species were positive for H5N5. Further, the dynamics of infection both to, and between the seals is unclear. Sampling of the seals was reactive to the avian die-off and so could have been missed as the mortality levels seen weren’t enough to trigger concern for seals at this heavily populated breeding site, and as such didn’t indicate a risk for HPAIV infection in seals. Environmental samples all tested negative suggesting that environmental contamination was low although in a dynamic and aggressive littoral environment coastal environment, both carcasses and environmentally relevant material (e.g., faecal material) may have been lost.

Recent studies have suggested that mammal-to-mammal transmission has occurred with H5N1 genotype B3.2 viruses in South America where significant mortality events have been seen [9-13]. The mortality level at Blakeney Point does not support that there was any mammal-to-mammal infection occurring, although the potential for this cannot be ruled out. Following the event, conservation groups involved with monitoring and assessing seal populations around the British Isles were notified to be on increased alert in case the infection continued in pinnipeds population but no further cases of H5N5 in seals was detected in GB or reported from Europe. Grey seals in GB are part of a pan-European metapopulation and exhibit partial migration [55]. Interestingly, one female seal (cow), which was found dead in this incident had been tagged at the Ecomare seal centre on the Dutch Wadden Sea Island of Texel on the 29th of December 2014 as a mother-dependent whitecoat pup (pers. comm. Seal Centre Ecomare). This seal was eleven years old when it died. A higher proportion of the European metapopulation spend the breeding season in the UK compared to the rest of the year. As such, a substantial proportion of seals that breed at Blakeney Point likely subsequently travelled to continental Europe, particularly the Wadden Sea [56]. Even those remaining in Southeast England likely moved between haul-out sites (locations on land where seals come ashore to rest, breed, or give birth) [57]. As well as travelling offshore, Grey Seal foraging sites extend up to 450 km from a haul-out sites [58], and pups typically disperse more widely than adults [59]. Such movements mean that in the advent of mammal-to-mammal transmission, infection could spread widely through the North-east Atlantic Grey Seal population. Moreover, Grey Seals often haul-out in the vicinity of harbour seals. Indeed, Blakeney Point is part of The Wash & North Norfolk Special Area of Conservation (SAC), designated for Harbour Seals (the other seal species present in GB and of conservation concern). The SAC holds most of the English harbour seal population and has recently undergone unexplained declines. Grey seals have previously played a role in spreading disease (Phocine Distemper Virus) which has had a huge impact on the less wide-ranging Harbour Seal [60].

Infection of seals must therefore be likely attributed to close interactions with infected seabirds. These densely packed and dynamic seal colonies attract a regular traffic of wild birds to scavenge moribund and dead seals and in turn seals may scavenge infective wild birds which succumb to disease or become exposed to HPAIV from infective guano left by foraging birds. It is possible that such dense living conditions, along with likely exposure to faeces from local seabirds like gulls (as well as the scavenging of afterbirths and dead seal pups by gulls [26, 27] provides a likely interface for infection with HPAIV, especially through the winter season of maximal HPAIV infection pressure produced by its circulation in wild birds. Ingestion of seabirds by seals on the breeding colony is considered unlikely. With a high juvenile population and gull mortality events occurring in and around seals, it is more plausible that infection through interactions such as playing with carcasses or inhalation of infectious material during investigative behaviours around bird carcasses is responsible, although this cannot be conclusively proven.

The detection of the H5N1 subtype in a subset of sampled gulls at this mortality event may simply reflect overlapping ranges for gull populations. A further possibility is the scavenging of other wild birds by gulls within this population. Proximal to this event, a handful of species have tested positive for HPAIV following being reported to Defra via the wild bird surveillance initiative. In winter gulls are generally considered to be voracious opportunistic predators and scavengers and as such it is possible that the small number of gulls detected with H5N1 infection are the results of local infection through consumption of carrion.

A further interesting point is that all H5N5 sequence derived from the seals and gulls contained both the NA stalk deletion and in the E627K mutation associated with mammalian adaptation [61], further raising uncertainty on the directionality of infection. The most recently detected H5N5 sequences from different avian species included the stalk deletion but had a 627E residue at that position in PB2 [62]. However, assessment of global H5N5 sequence from avian species across the globe from 2020 onwards have demonstrated that the E627K mutation can be detected in avian species in the absence of obvious mammalian involvement [6, 7, 62]. The ability of birds to maintain viruses with mammalian adaptations is a further area of considerable interest, as where wild animal mortality events occur, omnivorous species are often found to be scavenging on mammalian carcasses which in turn gives the opportunity for viruses that are already partially adapted for mammalian infection to circulate in wild birds with a potentially heightened risk of mammalian infection if further spill over into terrestrial mammals occurs [9-13]. This phenomenon requires further investigation.

The co-circulation of distinct subtypes and genotypes within species occupying the same temporal, spatial and ecological landscapes raise important questions regarding the potential for viral reassortment and the role of gulls in the emergence of novel genotypes. Numerous H5N1 genotypes have been described in each previous season dominated by H5N1 viruses, although a few genotypes were detected more frequently [35, 54]. Interestingly, since the beginning of the 2024/25 AIV season in GB, genotype diversity has been limited with DI.2 dominating positive avian detections [62]. Alongside DI.2, only the BB genotype, a previously dominant genotype, has been detected with some regularity, predominantly in the South-West of England in gull species that links with occasional detections in continental Europe [54]. Further, there have been two detections of the DI.1 genotype in wild birds in 2024, both have been of Norfolk, and it has not been detected since. Interestingly the contemporary H5N5 detections appear to be entirely represented by a single genotype (genotype I), with no evidence of reassortment. Further, since the emergence of the H5N1 BB genotype in 2022, this has also exhibited predominately in gull species with limited reassortment. How and why different HPAIV genotypes are generated, and either dominate or disappear, remains a significant knowledge gap although several studies have demonstrated differential shedding and clinical impact in poultry species and it is likely that similar occurs in wild birds.

In conclusion, the detection of two subtypes, H5N1 and H5N5 in avian species linked to a mammalian mortality event that appears restricted to infection with a single H5N5 subtype is of high interest. Conditions for sampling carcasses were sub-optimal and the environmental conditions within which these detections were made precluded further swabbing and assessment. Regardless, it is important to define and describe such detections as they clearly demonstrate that different viral subtypes can be present within a mortality event without reassortment being detected. Understanding the intra and inter-species infection and transmission dynamics is of high interest.

## Acknowledgements

Field work to undertake extended sampling of seal and avian carcasses, as well as testing and virus characterisation was supported by the Defra funded iPREPARE initiative (SE2230). Broader testing and generation of the viral sequences was funded by the Department for Environment, Food and Rural Affairs (Defra, UK) and the Devolved Administrations of Scotland and Wales, through the following grants: SV3032, SV3400, SE2227 and OMO402. This work was also supported by the Biotechnology and Biological Sciences Research Council (BBSRC) and Department for Environment, Food and Rural Affairs (Defra, UK) research initiative ‘FluTrailMap’ [grant number BB/Y007271/1]. Funded by the European Union under grant agreement (101084171) - (Kappa-Flu). Views and opinions expressed are, however, those of the author(s) only and do not necessarily reflect those of the European Union or REA. Neither the European Union nor the granting authority can be held responsible for them. We acknowledge the National Trust for rapid reporting of avian mortality via the Defra online reporting system and for enabling access to the site by those involved in sampling. DJFR’s contribution was funded by NERC National Capability Funding (NE/Y006194/1). DV was supported by a UKRI fellowship (BB/W009404/1).

## Figure legends

**Supplementary Figure 1:** Maximum likelihood phylogeny of European H5 Influenza sequences subset to represent 0.5% sequence divergence. Tips are coloured by region of sequence submission and tip shapes indicate whether a sequence was in the GISAID database or generated for this study.

## References

1. Krammer, F., E. Hermann, and A.L. Rasmussen, Highly pathogenic avian influenza H5N1: history, current situation, and outlook. Journal of Virology, 2025. 99(4): p. e02209–24.

2. Bellido-Martín, B., et al., Evolution, spread and impact of highly pathogenic H5 avian influenza A viruses. Nature Reviews Microbiology, 2025.

3. Caliendo, V., et al., Long-Term Protective Effect of Serial Infections with H5N8 Highly Pathogenic Avian Influenza Virus in Wild Ducks. J Virol, 2022. 96(18): p. e0123322.

4. Byrne, A.M.P., et al., Investigating the Genetic Diversity of H5 Avian Influenza Viruses in the United Kingdom from 2020–2022. Microbiology Spectrum, 2022. 11(4): p. e04776–22.

5. Peacock, T.P., et al., The global H5N1 influenza panzootic in mammals. Nature, 2025. 637(8045): p. 304–313.

6. European Food Safety Authority (EFSA), et al., Avian influenza overview December 2023– March 2024. Efsa Journal, 2024. 22(3): p. e8754.

7. Erdelyan, C.N.G., et al., Multiple transatlantic incursions of highly pathogenic avian influenza clade 2.3.4.4b A(H5N5) virus into North America and spillover to mammals. Cell Rep, 2024. 43(7): p. 114479.

8. Hew, Y.L., et al., Cocirculation of Genetically Distinct Highly Pathogenic Avian Influenza H5N5 and H5N1 Viruses in Crows, Hokkaido, Japan. Emerg Infect Dis, 2024. 30(9): p. 1912–1917.

9. Leguia, M., et al., Highly pathogenic avian influenza A (H5N1) in marine mammals and seabirds in Peru. Nature Communications, 2023. 14(1): p. 5489.

10. Pardo-Roa, C., et al., Cross-species and mammal-to-mammal transmission of clade 2.3.4.4b highly pathogenic avian influenza A/H5N1 with PB2 adaptations. Nature Communications, 2025. 16(1): p. 2232.

11. Uhart, M.M., et al., Epidemiological data of an influenza A/H5N1 outbreak in elephant seals in Argentina indicates mammal-to-mammal transmission. Nature Communications, 2024. 15(1): p. 9516.

12. Banyard, A.C., et al., Detection and spread of high pathogenicity avian influenza virus H5N1 in the Antarctic Region. Nat Commun, 2024. 15(1): p. 7433.

13. Bennison, A., et al., A case study of highly pathogenic avian influenza (HPAI) H5N1 at Bird Island, South Georgia: the first documented outbreak in the subantarctic region. Bird Study, 2024. 71(4): p. 380–391.

14. Agüero, M., et al., Highly pathogenic avian influenza A(H5N1) virus infection in farmed minks, Spain, October 2022. Eurosurveillance, 2023. 28(3): p. 2300001.

15. Kareinen, L., et al., Highly pathogenic avian influenza A(H5N1) virus infections on fur farms connected to mass mortalities of black-headed gulls, Finland, July to October 2023. Euro Surveill, 2024. 29(25).

16. Banyard, A.C., et al., Detection of clade 2.3.4.4b H5N1 high pathogenicity avian influenza virus in a sheep in Great Britain, 2025. bioRxiv, 2025: p. 2025.06.27.661969.

17. USA Centers for Disease Control and Prevention (CDC). H5 Bird Flu: Current Situation. 2025 [cited 2025 April]; Available from: https://www.cdc.gov/bird-flu/situation-summary/index.html.

18. Caserta, L.C., et al., Spillover of highly pathogenic avian influenza H5N1 virus to dairy cattle. Nature, 2024. 634(8034): p. 669–676.

19. Garg, S., et al., Highly Pathogenic Avian Influenza A(H5N1) Virus Infections in Humans. N Engl J Med, 2025. 392(9): p. 843–854.

20. Spackman, E., et al., Characterization of highly pathogenic avian influenza virus in retail dairy products in the US. J Virol, 2024. 98(7): p. e0088124.

21. Falchieri, M., et al., Rapid mortality in captive bush dogs (Speothos venaticus) caused by influenza A of avian origin (H5N1) at a wildlife collection in the United Kingdom. Emerg Microbes Infect, 2024. 13(1): p. 2361792.

22. Naraharisetti, R., et al., Highly Pathogenic Avian Influenza A(H5N1) Virus Infection of Indoor Domestic Cats Within Dairy Industry Worker Households - Michigan, May 2024. MMWR Morb Mortal Wkly Rep, 2025. 74(5): p. 61–65.

23. The Animal and Plant Health Agency (APHA) and The Department for Environment Food and Rural Affairs (DEFRA). Confirmed findings of influenza of avian origin in non-avian wildlife. 2025 [cited 2025 April]; Available from: https://www.gov.uk/government/publications/bird-flu-avian-influenza-findings-in-non-avian-wildlife/confirmed-findings-of-influenza-of-avian-origin-in-non-avian-wildlife.

24. The National Trust. Blakeney National Nature Reserve. 2025 [cited 2025 June]; Available from: https://www.nationaltrust.org.uk/visit/norfolk/blakeney-national-nature-reserve.

25. Morris, C., et al., List of Briefing Papers for SCOS. Scientific Advice on Matters Related to the Management of Seal Populations: 2024: p. 123.

26. Twiss, S.D., C. Duck, and P.P. Pomeroy, Grey seal (Halichoerus grypus) pup mortality not explained by local breeding density on North Rona, Scotland. Journal of Zoology, 2003. 259(1): p. 83–91.

27. Quaggiotto, M.-M., et al., Seal carrion is a predictable resource for coastal ecosystems. Acta Oecologica, 2018. 88: p. 41–51.

28. The Department for Food and Rural Affairs (DEFRA) and The Animal and Plant Health Agency (APHA). Guidance: Report dead wild birds. 2025 [cited 2025 June]; Available from: https://www.gov.uk/guidance/report-dead-wild-birds.

29. The Department for Food and Rural Affairs (DEFRA) and The Animal and Plant Health Agency (APHA). Guidance: Influenza A (H5N1) infection in mammals: suspect case definition and diagnostic testing criteria. 2025 [cited 2025 June]; Available from: https://www.gov.uk/government/publications/listed-diseases-in-animals-case-definitions-testing-and-reporting/influenza-a-h5n1-infection-in-mammals-suspect-case-definition-and-diagnostic-testing-criteria.

30. James, J., et al., The Role of Airborne Particles in the Epidemiology of Clade 2.3.4.4b H5N1 High Pathogenicity Avian Influenza Virus in Commercial Poultry Production Units. Viruses, 2023. 15(4).

31. James, J., et al., Clade 2.3.4.4b H5N1 high pathogenicity avian influenza virus (HPAIV) from the 2021/22 epizootic is highly duck adapted and poorly adapted to chickens. J Gen Virol, 2023. 104(5).

32. Nagy, A., et al., A universal RT-qPCR assay for “One Health” detection of influenza A viruses. PLoS One, 2021. 16(1): p. e0244669.

33. James, J., et al., Rapid and sensitive detection of high pathogenicity Eurasian clade 2.3.4.4b avian influenza viruses in wild birds and poultry. J Virol Methods, 2022. 301: p. 114454.

34. James, J., et al., Development and Application of Real-Time PCR Assays for Specific Detection of Contemporary Avian Influenza Virus Subtypes N5, N6, N7, N8, and N9. Avian Dis, 2019. 63(p1): p. 209–218.

35. Byrne, A.M.P., et al., Investigating the Genetic Diversity of H5 Avian Influenza Viruses in the United Kingdom from 2020–2022. Microbiology Spectrum, 2023. 11(4): p. e04776–22.

36. Katoh, K. and D.M. Standley, MAFFT multiple sequence alignment software version 7: improvements in performance and usability. Molecular biology and evolution, 2013. 30(4): p. 772–780.

37. Shen, W., B. Sipos, and L. Zhao, SeqKit2: A Swiss army knife for sequence and alignment processing. iMeta, 2024. 3(3): p. e191.

38. Minh, B.Q., et al., IQ-TREE 2: new models and efficient methods for phylogenetic inference in the genomic era. Molecular biology and evolution, 2020. 37(5): p. 1530–1534.

39. Kalyaanamoorthy, S., et al., ModelFinder: fast model selection for accurate phylogenetic estimates. Nature methods, 2017. 14(6): p. 587–589.

40. Hoang, D.T., et al., UFBoot2: improving the ultrafast bootstrap approximation. Molecular biology and evolution, 2018. 35(2): p. 518–522.

41. Sagulenko, P., V. Puller, and R.A. Neher, TreeTime: Maximum-likelihood phylodynamic analysis. Virus evolution, 2018. 4(1): p. vex042.

42. Wickham, H., ggplot2: Elegant Graphics for Data Analysis. 2016, New York: Springer-Verlag

43. Guangchuang, Y.D., K., Smith; Huachen, Zhu; Yi Guan; Tommy, Tsan-Yuk, Lam, ggtree: an r package for visualization and annotation of phylogenetic trees with their covariates and other associated data. Methods Ecol Evol, 2017. 8: p. 28–36.

44. Larsson, A., AliView: a fast and lightweight alignment viewer and editor for large datasets. Bioinformatics, 2014. 30(22): p. 3276–3278.

45. Reid, S.M., et al., A multi-species, multi-pathogen avian viral disease outbreak event: Investigating potential for virus transmission at the wild bird – poultry interface. Emerging Microbes & Infections, 2024. 13(1): p. 2348521.

46. Ross, C.S., et al., Genetic Analysis of H5N1 High-Pathogenicity Avian Influenza Virus following a Mass Mortality Event in Wild Geese on the Solway Firth. Pathogens, 2024. 13(1).

47. Banyard, A.C., et al., Detection of Highly Pathogenic Avian Influenza Virus H5N1 Clade 2.3.4.4b in Great Skuas: A Species of Conservation Concern in Great Britain. Viruses, 2022. 14(2): p. 212.

48. Falchieri, M., et al., Shift in HPAI infection dynamics causes significant losses in seabird populations across Great Britain. Vet Rec, 2022. 191(7): p. 294–296.

49. Floyd, T., et al., Encephalitis and Death in Wild Mammals at a Rehabilitation Center after Infection with Highly Pathogenic Avian Influenza A(H5N8) Virus, United Kingdom. Emerg Infect Dis, 2021. 27(11): p. 2856–2863.

50. Fusaro, A., et al., High pathogenic avian influenza A(H5) viruses of clade 2.3.4.4b in Europe— Why trends of virus evolution are more difficult to predict. Virus Evolution, 2024. 10(1).

51. Burrough, E.R., et al., Highly Pathogenic Avian Influenza A(H5N1) Clade 2.3.4.4b Virus Infection in Domestic Dairy Cattle and Cats, United States, 2024. Emerg Infect Dis, 2024. 30(7): p. 1335–1343.

52. Nguyen, T.-Q., et al., Emergence and interstate spread of highly pathogenic avian influenza A(H5N1) in dairy cattle in the United States. Science, 2025. 388(6745): p. eadq0900.

53. The Animal and Plant Health Agency (APHA). Research and analysis: Bird flu (avian influenza): cases in wild birds. 2025 [cited 2025 Jan]; Available from: https://www.gov.uk/government/publications/avian-influenza-in-wild-birds.

54. Alexakis, L., et al., Avian influenza overview December 2024-March 2025. Efsa j, 2025. 23(4): p. e9352.

55. Russell, D.J.F., et al., Uncovering the links between foraging and breeding regions in a highly mobile mammal. Journal of Applied Ecology, 2013. 50(2): p. 499–509.

56. Brasseur, S.M.J.M., et al., Rapid recovery of Dutch gray seal colonies fueled by immigration. Marine Mammal Science, 2015. 31(2): p. 405–426.

57. Russell, D., Movements of grey seal that haul out on the UK coast of the southern North Sea. Sea Mammal Research Unit Report to the Department of Energy and Climate Change (OESEA-14-17), 2016: p. 18.

58. Carter, M.I.D., et al., Sympatric Seals, Satellite Tracking and Protected Areas: Habitat-Based Distribution Estimates for Conservation and Management. Frontiers in Marine Science, 2022. Volume 9 - 2022.

59. Carter, M.I.D., et al., Intrinsic and extrinsic factors drive ontogeny of early-life at-sea behaviour in a marine top predator. Scientific Reports, 2017. 7(1): p. 15505.

60. Härkönen, T., et al., The 1988 and 2002 phocine distemper virus epidemics in European harbour seals. Dis Aquat Organ, 2006. 68(2): p. 115–30.

61. Long, J.S., et al., Species difference in ANP32A underlies influenza A virus polymerase host restriction. Nature, 2016. 529(7584): p. 101–104.

62. The Animal and Plant Health Agency (APHA) and The Department for Environment Food and Rural Affairs (DEFRA). Updated Outbreak Assessment #7: High pathogenicity avian influenza (HPAI) in Great Britain and Europe. 2025 [cited 2025 June]; Available from: https://assets.publishing.service.gov.uk/media/67e158500114b0b86e59f4d9/HPAI_in_Grea t_Britain_and_Europe_March_2025.pdf.

63. Leyson, C., et al., Pathogenicity and genomic changes of a 2016 European H5N8 highly pathogenic avian influenza virus (clade 2.3.4.4) in experimentally infected mallards and chickens. Virology, 2019. 537: p. 172–185.

64. Chrzastek, K., et al., Use of Sequence-Independent, Single-Primer-Amplification (SISPA) for rapid detection, identification, and characterization of avian RNA viruses. Virology, 2017. 509: p. 159–166.

