## Supplementary tables for "Co-circulation of distinct high pathogenicity avian influenza virus (HPAIV) subtypes in a mass mortality event in wild seabirds and co-location with dead seals"

### Supplementary Materials

**Supplementary Table 1:** Oligonucleotide primers used for cDNA synthesis and amplification for whole genome sequencing.

| **Name** | **Sequence** | **Reference** |
| --- | --- | --- |
| Optil – F1 | TTACGCGCCAGCAAAAGCAG | [63]; [64] |
| Optil – F2 | GTTACGCGCCAGCGAAAGCAGG |  |
| Optil – R1 | GTTACGCGCCAGTAGAAACAAG |  |

**Supplementary Table 2:** GISAID Accession numbers for all sequences generated in this study.

| **Common Name** | **Sample** | **Collection Date** | **Subtype (genotype)** | **Virus Strain Name** | **GISAID accession number** |
| --- | --- | --- | --- | --- | --- |
| GBBG | Brain | 2025-02-14 | H5N1 (DI.2) | A/Greater_Black_Backed_Gull/England/014814/2025 | EPI_ISL_19779797 |
| Grey seal | Rectal | 2025-02-14 | H5N5 (I) | A/Grey_Seal/England/014816/2025 | EPI_ISL_19779798 |
| GBBG | Cloacal Swab | 2025-02-14 | H5N5 (I) | A/Greater_Black_Backed_Gull/England/014827/2025 | EPI_ISL_19779799 |
| Herring gull | OP swab | 2025-02-14 | H5N5 (I) | A/Herring_Gull/England/014794/2025 | EPI_ISL_19779796 |
| Grey seal | Brain | 2025-02-14 | H5N5 (I) | A/Grey_Seal/England/014989/2025 | EPI_ISL_19779804 |
| Grey seal | Brain | 2025-02-14 | H5N5 (I) | A/Grey_Seal/England/014976/2025 | EPI_ISL_19779803 |
| Grey seal | Brain | 2025-02-14 | H5N5 (I) | A/Grey_Seal/England/014941/2025 | EPI_ISL_19779802 |
| Grey seal | Brain | 2025-02-14 | H5N5 (I) | A/Grey_Seal/England/014917/2025 | EPI_ISL_19779801 |
| Grey seal | Brain | 2025-02-14 | H5N5 (I) | A/Grey_Seal/England/014913/2025 | EPI_ISL_19779800 |
| Northern Fulmar | Brain | 2025-02-14 | H5N5 (I) | A/Fulmar/England/016501/2025 | EPI_ISL_19848222 |
| Northern Fulmar | Cloacal swab | 2025-02-14 | H5N5 (I) | A/Fulmar/England/016619/2025 | EPI_ISL_19848173 |
| Northern Fulmar | OP swab | 2025-02-14 | H5N5 (I) | A/Fulmar/England/016621/2025 | EPI_ISL_19848174 |

GBBG = Great Black-backed Gull

**Supplementary table 3:** Samples collected and results.

| Preliminary first investigation | | | | | | | | |
| --- | --- | --- | --- | --- | --- | --- | --- | --- |
| **Sample number** | **Species** | **Animal number** | **Sampled tissue / organ** | **Nagy (M)** | **HP-H5** | **N1** | **N5** | **Interpretation** |
| 1 | Grey Seal | 1 | Oral | NEG | NEG | NEG | NEG | Neg |
| 2 |  |  | Rectal | NEG | NEG | NEG | NEG |  |
| 3 |  |  | Brain | NEG | NEG | NEG | NEG |  |
| 4 | Grey Seal | 2 | Oral | NEG | NEG | NEG | NEG | Neg |
| 5 |  |  | Rectal | NEG | NEG | NEG | NEG |  |
| 6 |  |  | Brain | NEG | NEG | NEG | NEG |  |
| 7 | Grey Seal | 3 | Oral | **POS** | **POS** | NEG | **POS** | HP H5N5 |
| 8 |  |  | Rectal | NEG | **POS** | NEG | **POS** |  |
| 9 |  |  | Brain | **POS** | **POS** | NEG | **POS** |  |
| 10 | Grey Seal | 4 | Oral | **POS** | **POS** | NEG | **POS** | HP H5N5 |
| 11 |  |  | Rectal | **POS** | **POS** | NEG | **POS** |  |
| 12 |  |  | Brain | **POS** | **POS** | NEG | **POS** |  |
| 13 | Herring Gull | 5 | OP | NEG | NEG | NEG | NEG | NEG |
| 14 |  |  | Cloacal | NEG | NEG | NEG | NEG |  |
| 15 | GBBG | 6 | OP | **POS** | **POS** | **POS** | NEG | HP H5N1 |
| 16 |  |  | Cloacal | **POS** | **POS** | **POS** | NEG |  |
| 17 | GBBG | 7 | OP | NEG | NEG | NEG | NEG | NEG |
| 18 |  |  | Cloacal | NEG | NEG | NEG | NEG |  |
| 19 | GBBG | 8 | OP | **POS** | **POS** | NEG | **POS** | HP H5N5 |
| 20 |  |  | Cloacal | **POS** | **POS** | NEG | **POS** |  |
| 21 | GBBG | 9 | OP | **POS** | **POS** | **POS** | NEG | HP H5N1 |
| 22 |  |  | Cloacal | **POS** | **POS** | **POS** | NEG |  |
| Second investigation | | | | | | | | |
| Sample number | Species | Animal number | Sampled tissue / organ | Nagy (M) | HPH5 | N1 | N5 | Conclusion |
| 1 | Atlantic Puffin | 1 | Brain | ND | **POS** | NEG | **POS** | HP H5N5 |
| 2 |  |  | Lung | ND | **POS** | NEG | **POS** |  |
| 3 |  |  | Trachea | ND | **POS** | NEG | **POS** |  |
| 4 |  |  | Heart | ND | **POS** | NEG | **POS** |  |
| 5 |  |  | Liver | ND | **POS** | NEG | **POS** |  |
| 6 |  |  | Spleen | ND | **POS** | NEG | **POS** |  |
| 7 |  |  | Kidney | ND | **POS** | NEG | **POS** |  |
| 8 |  |  | Jejunum | ND | **POS** | NEG | **POS** |  |
| 9 |  |  | Oviduct | ND | **POS** | NEG | **POS** |  |
| 10 |  |  | Cloacal | ND | NEG | NEG | NEG |  |
| 11 |  |  | OP | ND | NEG | NEG | NEG |  |
| 12 | Northern Fulmar | 2 | Brain | ND | **POS** | NEG | **POS** | HP H5N5 |
| 13 |  |  | Lung | ND | **POS** | NEG | **POS** |  |
| 14 |  |  | Trachea | ND | **POS** | NEG | **POS** |  |
| 15 |  |  | Heart | ND | **POS** | NEG | **POS** |  |
| 16 |  |  | Liver | ND | **POS** | NEG | **POS** |  |
| 17 |  |  | Spleen | ND | **POS** | NEG | **POS** |  |
| 18 |  |  | Kidney | ND | **POS** | NEG | **POS** |  |
| 19 |  |  | Jejunum | ND | **POS** | NEG | **POS** |  |
| 20 |  |  | Oviduct | ND | **POS** | NEG | **POS** |  |
| 21 |  |  | Cloacal | ND | **POS** | NEG | **POS** |  |
| 22 |  |  | OP | ND | **POS** | NEG | **POS** |  |
| 23 | GBBG | 3 | Brain | NEG | **POS** | NEG | **POS** | HP H5N5 |
| 24 | Grey Seal | 4 | Nasal | NEG | NEG | NEG | NEG | NEG |
| 25 |  |  | Brain | NEG | NEG | NEG | NEG |  |
| 26 |  |  | Rectal | NEG | NEG | NEG | NEG |  |
| 27 | Grey Seal | 5 | Rectal | NEG | NEG | NEG | NEG | NEG |
| 28 |  |  | Nasal | NEG | NEG | NEG | NEG |  |
| 29 |  |  | Oral | NEG | NEG | NEG | NEG |  |
| 30 |  |  | Brain | NEG | NEG | NEG | NEG |  |
| 31 | GBBG | 6 | OP | **POS** | **POS** | NEG | **POS** | HP H5N5 |
| 32 | Herring Gull | 7 | OP | **POS** | **POS** | NEG | **POS** | HP H5N5 |
| 33 | Grey Seal | 8 | Rectal | NEG | NEG | NEG | NEG | NEG |
| 34 |  |  | Nasal | NEG | NEG | NEG | NEG |  |
| 35 |  |  | Oral | NEG | NEG | NEG | NEG |  |
| 36 |  |  | Brain | NEG | NEG | NEG | NEG |  |
| 37 | GBBG | 9 | OP | **POS** | **POS** | NEG | **POS** | HP H5N5 |
| 38 |  |  | Brain | **POS** | **POS** | NEG | **POS** |  |
| 39 | Grey Seal | 10 | Rectal | NEG | NEG | NEG | NEG | NEG |
| 40 | Grey Seal | 11 | Rectal | NEG | NEG | NEG | NEG | NEG |
| 41 | Grey Seal | 12 | Rectal | **POS** | **POS** | NEG | **POS** | HP H5N5 |
| 42 | Grey Seal | 13 | Rectal | NEG | NEG | NEG | NEG | NEG |
| 43 | Grey Seal | 14 | Rectal | NEG | NEG | NEG | NEG | NEG |
| 44 | Grey Seal | 15 | Rectal | NEG | NEG | NEG | NEG | NEG |
| 45 | Grey Seal | 16 | Rectal | NEG | NEG | NEG | NEG | NEG |
| 46 | Grey Seal | 17 | Rectal | NEG | NEG | NEG | NEG | NEG |
| 47 | GBBG | 18 | OP | **POS** | **POS** | NEG | **POS** | HP H5N5 |
| 48 | GBBG | 19 | OP | **POS** | **POS** | **POS** | NEG | HP H5N1 |
| 49 | GBBG | 20 | OP | **POS** | **POS** | **POS** | NEG | HP H5N1 |
| 50 |  |  | Brain | **POS** | **POS** | **POS** | NEG |  |
| 51 |  |  | Cloacal | NEG | NEG | NEG | NEG |  |
| 52 | Grey Seal | 21 | Rectal | **POS** | **POS** | NEG | **POS** | HP H5N5 |
| 53 | Grey Seal | 22 | Rectal | **POS** | **POS** | NEG | **POS** | HP H5N5 |
| 54 | Grey Seal | 23 | Rectal | **POS** | **POS** | NEG | **POS** | HP H5N5 |
| 55 | GBBG | 24 | OP | **POS** | **POS** | NEG | **POS** | HP H5N5 |
| 56 | Grey Seal | 25 | Oral | NEG | NEG | NEG | NEG | NEG |
| 57 |  |  | Rectal | NEG | NEG | NEG | NEG |  |
| 58 | GBBG | 26 | OP | **POS** | **POS** | NEG | **POS** | HP H5N5 |
| 59 | Northern Fulmar | 27 | OP | **POS** | **POS** | **POS** | NEG | HP H5N1 |
| 60 | Grey Seal | 28 | Nasal | **POS** | **POS** | NEG | **POS** | HP H5N5 |
| 61 | GBBG | 29 | OP | NEG | NEG | NEG | NEG | NEG |
| 62 | GBBG | 30 | OP | **POS** | **POS** | NEG | **POS** | HP H5N5 |
| 63 |  |  | Cloacal | **POS** | **POS** | NEG | **POS** |  |
| 64 | Grey Seal | 31 | Rectal | NEG | NEG | NEG | NEG | NEG |
| 65 | Grey Seal | 32 | Nasal | **POS** | NEG | NEG | NEG | HP H5N5 |
| 66 |  |  | Oral | **POS** | **POS** | NEG | **POS** |  |
| 67 |  |  | Rectal | **POS** | **POS** | NEG | **POS** |  |
| 68 |  |  | Brain | **POS** | **POS** | NEG | **POS** |  |
| 69 |  |  | Lung | NEG | NEG | NEG | NEG |  |
| 70 | GBBG | 33 | OP | NEG | NEG | NEG | NEG | NEG |
| 71 |  |  | Cloacal | NEG | NEG | NEG | NEG |  |
| 72 | GBBG | 34 | OP | **POS** | **POS** | NEG | **POS** | HP H5N5 |
| 73 |  |  | Cloacal | **POS** | **POS** | NEG | **POS** |  |
| 74 | Grey Seal | 35 | Rectal | NEG | NEG | NEG | NEG | NEG |
| 75 | GBBG | 36 | OP | **POS** | **POS** | NEG | **POS** | HP H5N5 |
| 76 |  |  | Cloacal | **POS** | **POS** | NEG | **POS** |  |
| 77 | GBBG | 37 | OP | **POS** | **POS** | NEG | **POS** | HP H5N5 |
| 78 | GBBG | 38 | OP | **POS** | **POS** | NEG | **POS** | HP H5N5 |
| 79 | Grey Seal | 39 | Rectal | NEG | NEG | NEG | NEG | NEG |
| 80 | Grey Seal | 40 | Rectal | **POS** | **POS** | NEG | **POS** | HP H5N5 |
| 81 | Grey Seal | 41 | Oral/Nasal | NEG | NEG | NEG | NEG | NEG |
| 82 |  |  | Rectal | NEG | NEG | NEG | NEG |  |
| 83 |  |  | Brain | NEG | NEG | NEG | NEG |  |
| 84 | Grey Seal | 42 | Nasal | NEG | NEG | NEG | NEG | NEG |
| 85 |  |  | Oral | NEG | NEG | NEG | NEG |  |
| 86 |  |  | Rectal | NEG | NEG | NEG | NEG |  |
| 87 |  |  | Brain | NEG | NEG | NEG | NEG |  |
| 88 |  |  | Lung | NEG | NEG | NEG | NEG |  |
| 89 | Grey Seal | 43 | Nasal | NEG | NEG | NEG | NEG | HP H5N5 |
| 90 |  |  | Rectal | **POS** | **POS** | NEG | **POS** |  |
| 91 |  |  | Oral | **POS** | **POS** | NEG | **POS** |  |
| 92 |  |  | Brain | **POS** | **POS** | NEG | **POS** |  |
| 93 | Grey Seal | 44 | Rectal | NEG | NEG | NEG | NEG | HP H5N5 |
| 94 |  |  | Nasal | **POS** | **POS** | NEG | **POS** |  |
| 95 |  |  | Oral | NEG | NEG | NEG | NEG |  |
| 96 |  |  | Brain | **POS** | **POS** | NEG | **POS** |  |
| 97 | Grey Seal | 45 | Rectal | NEG | NEG | NEG | **POS** | HP H5N5 |
| 98 |  |  | Nasal | NEG | NEG | NEG | NEG |  |
| 99 |  |  | Oral | NEG | NEG | NEG | NEG |  |
| 100 |  |  | Brain | **POS** | **POS** | NEG | **POS** |  |
| 101 | Herring Gull | 46 | Cloacal | NEG | NEG | NEG | NEG | NEG |
| 102 |  |  | OP | NEG | NEG | NEG | NEG |  |
| 103 |  |  | Brain | NEG | NEG | NEG | NEG |  |
| 104 |  |  | Trachea | NEG | NEG | NEG | NEG |  |
| 105 |  |  | Alimentary tract | NEG | NEG | NEG | NEG |  |
| 106 | GBBG | 47 | OP | **POS** | **POS** | NEG | **POS** | HP H5N5 |
| 107 |  |  | Brain | **POS** | **POS** | NEG | **POS** |  |
| 108 | Grey Seal | 48 | Rectal | **POS** | NEG | NEG | NEG | HP H5N5 |
| 109 |  |  | Nasal | NEG | NEG | NEG | NEG |  |
| 110 |  |  | Oral | **POS** | **POS** | NEG | **POS** |  |
| 111 | Grey Seal | 49 | Predated carcass swabs, wounds swab | **POS** | **POS** | NEG | **POS** | HP H5N5 |
| 112 | GBBG | 50 | Brain | **POS** | NEG | NEG | **POS** | HP H5N5 |
| 113 |  |  | OP | **POS** | **POS** | NEG | **POS** |  |
| 114 | GBBG | 51 | Brain | **POS** | **POS** | NEG | **POS** | HP H5N5 |
| 115 |  |  | OP | **POS** | **POS** | NEG | NEG |  |
| 116 | Grey Seal | 52 | Rectal | **POS** | **POS** | NEG | **POS** | HP H5N5 |
| 117 |  |  | Oral | **POS** | NEG | NEG | NEG |  |
| 118 |  |  | Nasal | **POS** | **POS** | NEG | **POS** |  |
| 119 |  |  | Brain | **POS** | **POS** | NEG | **POS** |  |
| 120 | Grey Seal | 53 | Rectal | NEG | NEG | NEG | NEG | NEG |
| 121 |  |  | Oral | NEG | NEG | NEG | NEG |  |
| 122 |  |  | Nasal | NEG | NEG | NEG | NEG |  |
| 123 |  |  | Brain | NEG | NEG | NEG | NEG |  |
| 124 | GBBG | 54 | OP | **POS** | **POS** | NEG | **POS** | HP H5N5 |
| 125 |  |  | Brain | **POS** | **POS** | NEG | **POS** |  |
| 126 | Grey Seal | 55 | Brain | NEG | NEG | NEG | NEG | NEG |
| 127 | Grey Seal | 56 | Brain | NEG | NEG | NEG | NEG | NEG |
| 128 | Grey Seal | 57 | Brain | NEG | NEG | NEG | NEG | NEG |
| 129 | Grey Seal | 58 | Brain | NEG | NEG | NEG | NEG | NEG |
| 130 | Grey Seal | 59 | Brain | NEG | NEG | NEG | NEG | NEG |
| 131 | GBBG | 60 | OP | **POS** | **POS** | NEG | **POS** | HP H5N5 |
| 132 |  |  | Cloacal | **POS** | **POS** | NEG | **POS** |  |
| 133 |  |  | Brain | **POS** | **POS** | NEG | **POS** |  |
| 134 |  |  | Gizzard + intestine | **POS** | **POS** | NEG | **POS** |  |
| 135 |  |  | Trachea | **POS** | **POS** | NEG | **POS** |  |
| 136 | Razorbill | 61 | Cloacal | NEG | NEG | NEG | NEG | NEG |
| 137 |  |  | OP | NEG | NEG | NEG | NEG |  |
| 138 |  |  | Brain | NEG | NEG | NEG | NEG |  |
| 139 |  |  | Trachea | NEG | NEG | NEG | NEG |  |
| 140 |  |  | Gizzard + intestine | NEG | NEG | NEG | NEG |  |
| 141 | Grey Seal | 62 | Brain | NEG | NEG | NEG | NEG | NEG |
| 142 | Grey Seal | 63 | Brain | NEG | NEG | NEG | NEG | NEG |
| 143 |  |  | Heart | NEG | NEG | NEG | NEG |  |
| 144 |  |  | Lung | NEG | NEG | NEG | NEG |  |
| 145 |  |  | Intestine | NEG | NEG | NEG | NEG |  |
| 146 |  |  | Liver | NEG | NEG | NEG | NEG |  |
| 147 | Grey Seal | 64 | Brain | **POS** | **POS** | NEG | **POS** | HP H5N5 |
| 148 |  |  | Oral | **POS** | **POS** | NEG | **POS** |  |
| 149 |  |  | Nasal | **POS** | **POS** | NEG | **POS** |  |
| 150 |  |  | Rectal | **POS** | **POS** | NEG | **POS** |  |
| 151 | Grey Seal | 65 | Nasal | NEG | NEG | NEG | NEG | NEG |
| 152 |  |  | Oral | NEG | NEG | NEG | NEG |  |
| 153 |  |  | Brain | NEG | NEG | NEG | NEG |  |
| 154 |  |  | Rectal | NEG | NEG | NEG | NEG |  |
| 155 | GBBG | 66 | OP | **POS** | **POS** | **POS** | NEG | HP H5N1 |
| 156 |  |  | Cloacal | NEG | NEG | NEG | NEG |  |
| 157 |  |  | Brain | NEG | NEG | NEG | NEG |  |
| 158 | Grey Seal | 67 | Oral | **POS** | NEG | NEG | NEG | HP H5N5 |
| 159 |  |  | Nasal | **POS** | **POS** | NEG | **POS** |  |
| 160 |  |  | Brain | **POS** | **POS** | NEG | **POS** |  |
| 161 |  |  | Rectal | **POS** | **POS** | NEG | NEG |  |
| 162 | Environmental | - | Puddle water | NEG | NEG | NEG | NEG | NEG |
| 163 | Environmental | - | Feathers | NEG | NEG | NEG | NEG | NEG |
| 164 | Environmental | - | Feathers | NEG | NEG | NEG | NEG | NEG |
| 165 | Environmental | - | Feathers | NEG | NEG | NEG | NEG | NEG |
| 166 | Environmental | - | Bird faeces | NEG | NEG | NEG | NEG | NEG |
| 167 | Environmental | - | Bird faeces | NEG | NEG | NEG | NEG | NEG |
| 168 | Environmental | - | Goose faeces | NEG | NEG | NEG | NEG | NEG |
| 169 | Environmental | - | Goose faeces | NEG | NEG | NEG | NEG | NEG |
| 170 | Environmental | - | Goose faeces | NEG | NEG | NEG | NEG | NEG |
| 171 | Environmental | - | Gull / Wader faeces | NEG | NEG | NEG | NEG | NEG |
| 172 | Environmental | - | Gull / Wader faeces | NEG | NEG | NEG | NEG | NEG |
| 173 | Environmental | - | Gull / Wader faeces | NEG | NEG | NEG | NEG | NEG |

GBBG = Great Black-backed Gull; POS = positive; NEG = negative.
