## Supplementary figures and images for "Co-circulation of distinct high pathogenicity avian influenza virus (HPAIV) subtypes in a mass mortality event in wild seabirds and co-location with dead seals"

### Supplementary figure 1

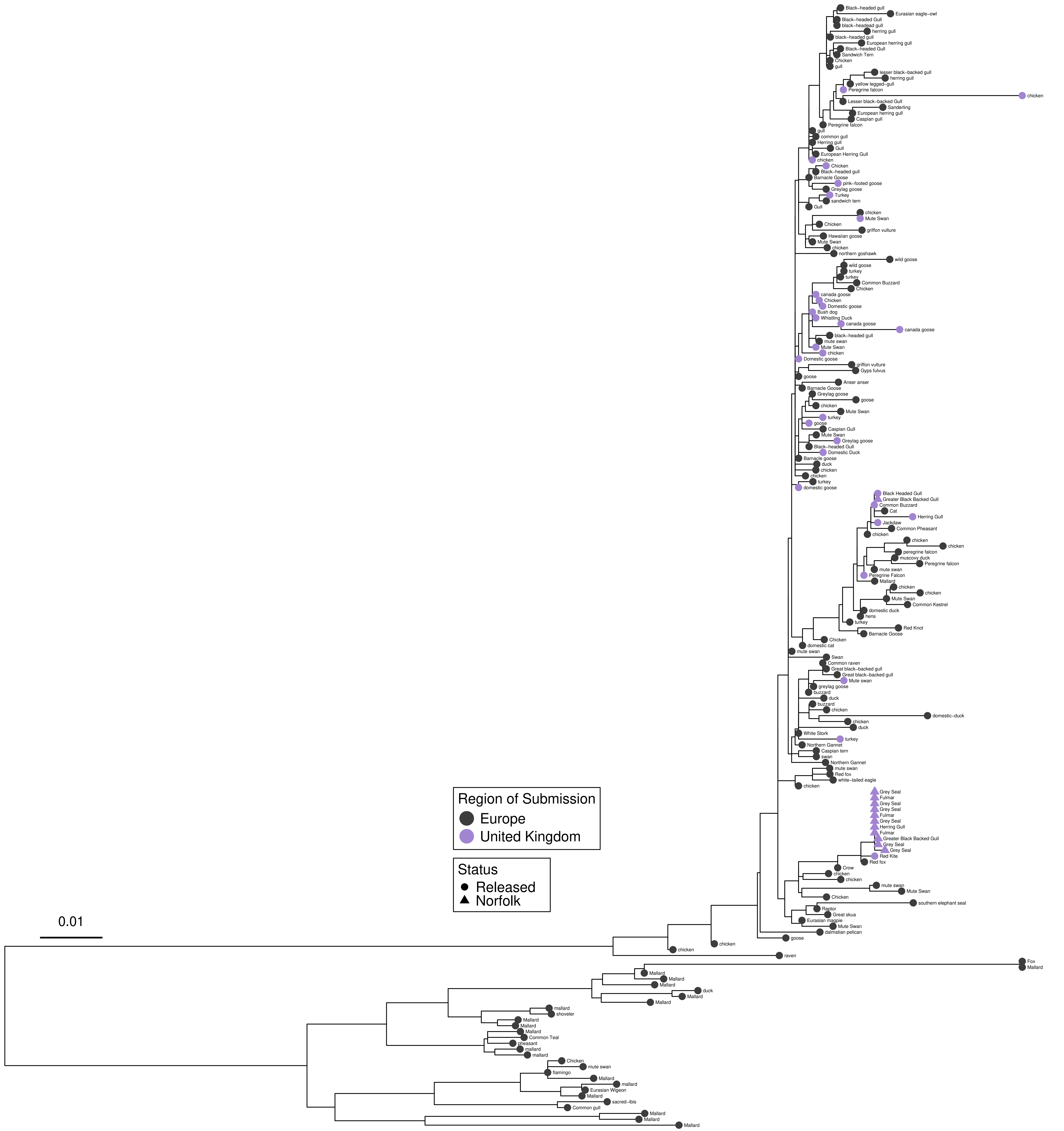
